# Image-based methods for phenotyping growth dynamics and fitness components in *Arabidopsis thaliana*

**DOI:** 10.1101/208512

**Authors:** François Vasseur, George Wang, Justine Bresson, Rebecca Schwab, Detlef Weigel

## Abstract

**Background:** The model species *Arabidopsis thaliana* has extensive resources to investigate intraspecific trait variability and the genetic bases of ecologically relevant traits. However, the cost of equipment and software required for high-throughput phenotyping is often a bottleneck for large-scale studies, such as mutant screening or quantitative genetics analyses. Simple tools are needed for the measurement of fitness-related traits, like relative growth rate and fruit production, without investment in expensive infrastructures. Here, we describe methods that enable the estimation of biomass accumulation and fruit number from the analysis of rosette and inflorescence images taken with a regular camera.

**Results:** We developed two models to predict plant dry mass and fruit number from the parameters extracted with the analysis of rosette and inflorescence images. Predictive models were trained by sacrificing growing individuals for dry mass estimation, and manually measuring a fraction of individuals for fruit number at maturity. Using a cross-validation approach, we showed that quantitative parameters extracted from image analysis predicts more 90% of both plant dry mass and fruit number. When used on 451 natural accessions, the method allowed modelling growth dynamics, including relative growth rate, throughout the life cycle of various ecotypes. Estimated growth-related traits had high heritability (0.65 < *H*^2^ < 0.93), as well as estimated fruit number (*H*^2^ = 0.68). In addition, we validated the method for estimating fruit number with *rev5*, a mutant with increased flower abortion.

**Conclusions:** The method we propose here is based on the automated computerization of plant images with ImageJ, and subsequent statistical modelling in R. It allows plant biologists to measure growth dynamics and fruit number in hundreds of individuals with simple computing steps that can be repeated and adjusted to a wide range of laboratory conditions. It is thus a flexible toolkit for the measurement of fitness-related traits in large plant populations.

## Background

Relative growth rate (RGR) and fruit number are two essential parameters of plant performance and fitness [1–3]. Proper estimation of RGR is achieved with the destructive measurement of plant biomass across several individuals sequentially harvested [4,5]. However, sequential harvesting is space and time consuming, which makes this approach inappropriate for large-scale studies. Furthermore, it is problematic for evaluating measurement error, as well as to compare growth dynamics and fitness-related traits, like fruit production, on the same individuals. Thus, a variety of platforms and equipment have been developed in the last decade for high-throughput phenotyping of plant growth from image analysis, specifically in crops [6–10] and in the model species *A. thaliana* [11–14]. Because commercial technologies are powerful but generally expensive [6,8,11,13], low-cost methods have been proposed, for instance to estimate rosette expansion rate from sequential imaging of *A. thaliana* individuals [14–16]. These methods can be adapted to a variety of lab conditions, but they do not allow the quantification of complex traits like biomass accumulation, RGR and fruit production.

Strong variation in RGR has been reported across and within plant species [17–22], which has been assumed to reflect the inherent diversity of strategies to cope with contrasting levels of resource availability [3,23,24]. For instance, species from resource-scarce environments generally show a lower RGR than species from resource-rich environments, even when they are grown in non-limiting resource conditions [25,26]. Ecophysiological studies [18,26] have shown that plant RGR depends on morphological traits (*e.g.* leaf mass fraction, leaf dry mass per area) and physiological rates (*e.g.* net assimilation rate) that differ between species, genotypes, or ontogenetic stages. For instance, plants become less efficient to accumulate biomass as they get larger and older, resulting in a decline of RGR during ontogeny [4]. This is due to developmental and allometric constraints such as self-shading and increasing allocation of biomass to supporting structures, like stems, in growing individuals.

To assess plant performance, response to environment, or genetic effects, it is important to link individual’s growth trajectory to productivity, yield or reproductive success. However, while several methods have been proposed to estimate growth dynamics from image analysis [8,11–16], methodologies for automated, high-throughput phenotyping of fruit number per plant remain surprisingly scarce [27,28]. Yet, the analysis of inflorescence images in *A. thaliana* could offer a valuable tool to connect growth dynamics and plant fitness. Because of its small size, inflorescence can easily be collected, imaged and analyzed with simple equipment. Furthermore, the genetic resources available in this species enable large-scale analyses (mutants screening, quantitative trait loci mapping and genome-wide association studies). For instance, the recent release of 1,135 natural accessions with complete genomic sequences allows conducting large comparative analysis of phenotypic variation within the species (http://1001genomes.org/) [29].

With the methods proposed here, we aimed at developing flexible and customizable tools based on the automated computerization and analysis of plant images to estimate fruit number and growth dynamics, including RGR throughout the life cycle, of many *A. thaliana* genotypes. The estimation of biomass accumulation is semi-invasive, as it requires sacrificing some individuals to train a predictive model. This approach considerably reduced the number of plants needed to estimate RGR during ontogeny, from seedling establishment to fruiting. Furthermore, the estimation of fruit number from automated image analysis of *A. thaliana* inflorescence could greatly help link growth variation to plant performance and fitness, in various genotypes and environmental conditions.

## Results

### Estimation of biomass accumulation, RGR and growth dynamics

#### Description

The method for growth analysis requires a set of rosette images on which we want to non-destructively measure plant dry mass, and a set of individuals harvested to train a predictive model (Fig. 1). In the case study presented here, we evaluated the method on 472 genotypes of *A. thaliana* grown in trays using a growth chamber equipped with Raspberry Pi Automated Phenotyping Array (hereafter RAPA) built at the Max Planck Institute (MPI) of Tübingen. We partitioned the whole population (*n* = 1920) in two subpopulations: the *focal* population (*n* = 960) on which growth dynamics (and fruit production) were measured, and the *training* population (*n* = 960) on which a predictive model of plant dry mass was developed.

**Figure 1.**
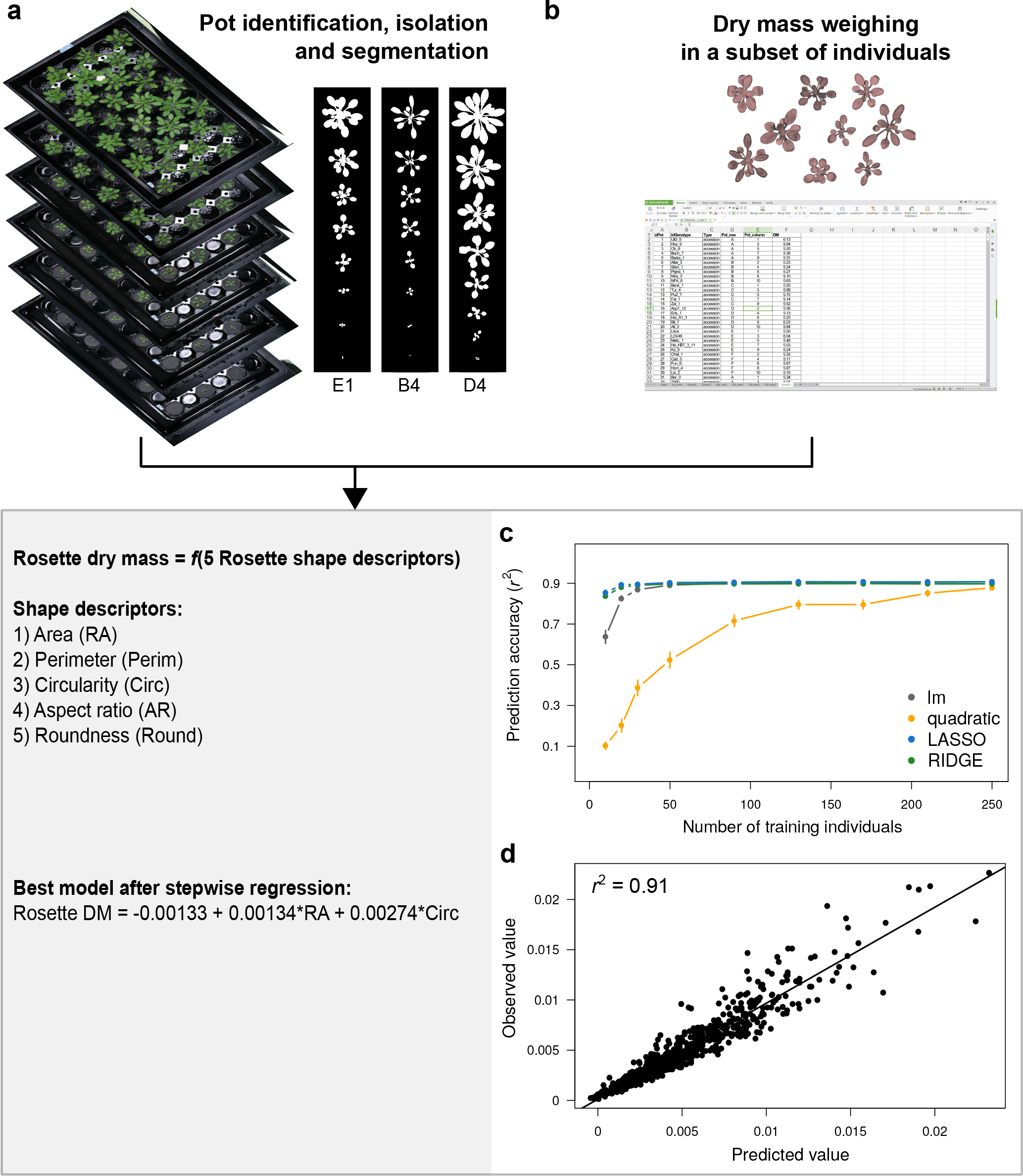
Estimation of plant dry mass from image analysis and statistical modelling. (**a**) Example of sequential tray images, analyzed with ImageJ to extract individual rosette shape descriptors during ontogeny. (**b**) Rosette dry mass measured at 16 DAG in the training population. (**c**) Series of cross-validation performed for different predictive models with different training population size (x axis). Dots represent mean prediction accuracy, measured as Pearson’s coefficient of correlation (*r*^2^) between observed and predicted values. Error bars represent 95% CI across 100 random permutations of the training dataset. (**d**) Correlation between observed and predicted values for cross-validation of the best model obtained with stepwise regression, performed 60 individuals to train the model, and tested on 300 individuals not used to train the model.

Individuals of the focal population were daily photographed during ontogeny (Fig. 1a), and harvested at the end of reproduction when the first fruits (siliques) were yellowing (stage 8.00 according to Boyes et al. [30]). Top-view images were automatically taken with RAPA, as well as with a manual setup during the first 25 days of plant growth (Fig. S1). To show the general applicability of the method, here we only used images manually taken with a regular camera. Moreover, the method can also be applied to images taken on individual pots. Plants of the training population were harvested at 16 days after germination (DAG), dried and weighed for building a predictive model of rosette biomass with top-view images (Fig. 1b). Predictive models were trained and evaluated with a cross-validation approach (Fig. 1c). Once a predictive model has been chosen and validated, rosette dry mass can be non-destructively estimated on all individuals of the focal population, which allows modelling growth trajectory, biomass accumulation and RGR throughout the plant life cycle.

#### Implementation

We developed an ImageJ [31] macro to extract shape descriptors of the rosette from tray or individual pot images (Fig, 1a, Additional File 1). To run the macro, users need to install ImageJ, go to *Plugins* > *Macros* > *Run*, and select the “RAPAmacro_RosetteShape.txt” file in the corresponding folder. Macro will guide users in the different steps of image analysis to label plant individual, perform segmentation and measure rosette shape descriptors. It proceeds all images (trays or individual pots) present in an input folder, and returns shape descriptors of individual rosettes in an output folder defined by users. Shape descriptors include individual rosette area and perimeter in pixels, rosette circularity (*Circ* = 4*π* × 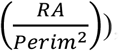), aspect ratio 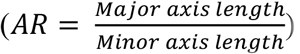, and roundness (*Round* = 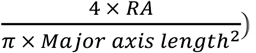. Rosette area and perimeter can be converted into cm^2^ and cm, respectively, by measuring the area and perimeter of a surface calibrator defined by users.

Predictive models of plant dry mass from shape descriptors were tested against measurements in the training population (Additional File 2). Depending on the training population size, we observed variable prediction accuracy for different models, as measured by the coefficient of correlation (*r*^2^) between measured and predicted rosette dry mass in individuals not used to train the model (Fig. 1c). LASSO and RIDGE models reached high prediction accuracy even with very small training population size (< 20 individuals). However, with a minimum of 50 training individuals, lm and RIDGE/LASSO performed equally, with a prediction accuracy > 90%. Using stepwise regression, we showed that using only rosette area (RA) and circularity (Circ) as predictors in a simple linear model framework can reach high prediction accuracy (*r*^2^ = 0.91, Fig. 1d). Thus, the final equation we used to estimate rosette dry mass from rosette pictures was *Rosette DM* = −0.00133 + 0.00134 × *RA* + 0.00274 × *Circ* (cross-validation *r*^2^ = 0.91, Fig. 1d).

#### Application

From estimated rosette dry mass during the ontogeny and final rosette dry mass measured on the same individuals at maturity, we modelled sigmoid growth curves of biomass accumulation (mg), *M*(t), for all individuals in the focal population with a three-parameter logistic function [4,32] (Fig. 2a and 2b), as in Equation 1:

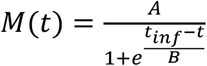

where *A*, *B* and *t*_inf_ are the parameters characterizing the shape of the curve, which differ between individuals depending on the genotypes and/or environmental conditions. *A* is the upper asymptote of the sigmoid curve, which was measured as rosette dry mass (mg) at maturity. The duration of growth was estimated as the time in days between the beginning of growth after vernalization (*t*_0_) and the end of reproduction (maturity). *B* controls the steepness of the curve, as the inverse of the exponential growth coefficient *r* (*r* = 1/*B*). *t*_inf_ is the inflection point that, by definition, corresponds to the point where the rosette is half the final dry mass. Both *B* and *t*_inf_ were estimated for every individual by fitting a logistic growth function to the data in R (Additional File 3).

**Figure 2.**
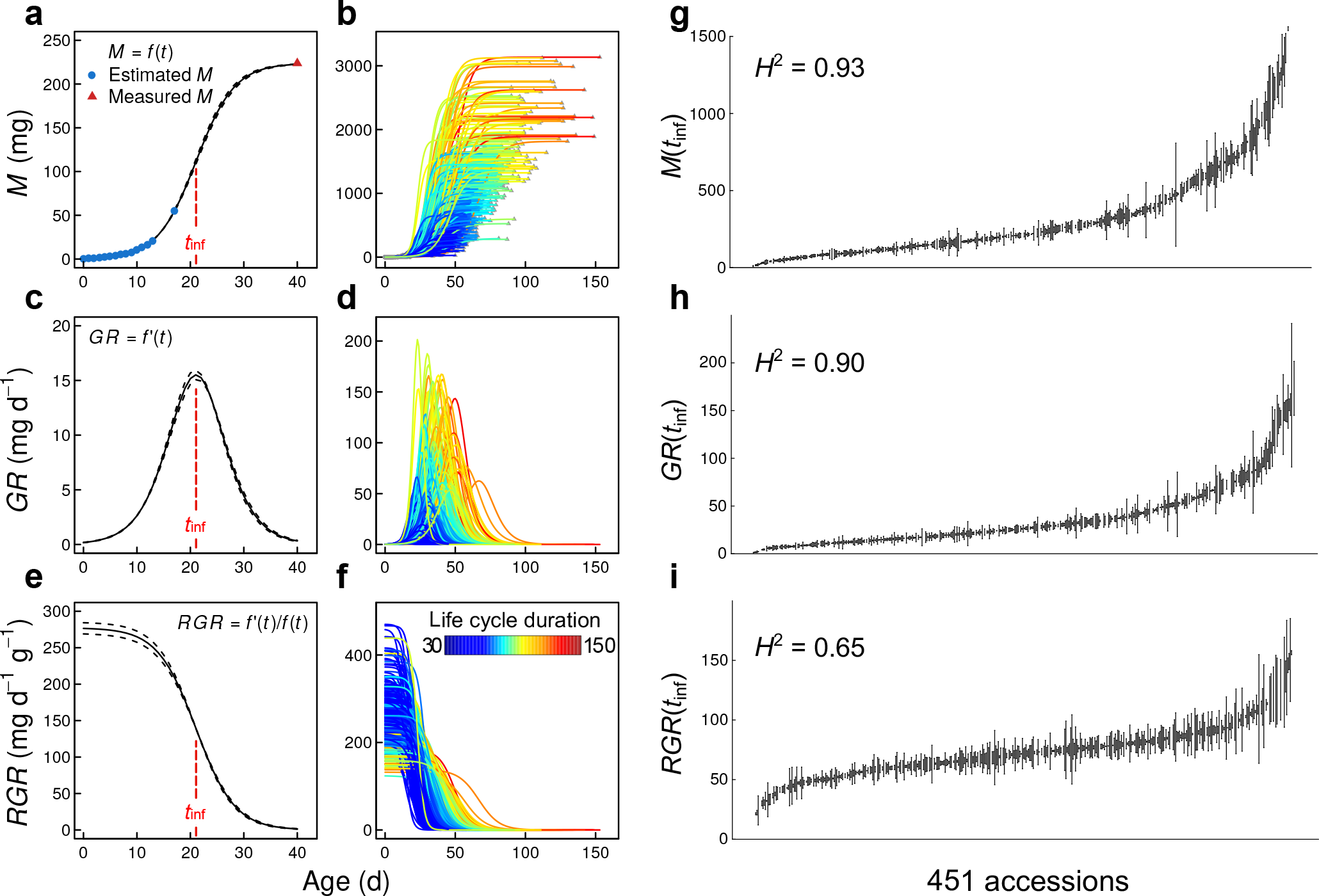
Application of the dry mass estimation method to model growth dynamics in *A. thaliana.* Statistical modelling of rosette dry mass during ontogeny, *M*(t), with three-parameter logistic growth curve, on one individual (**a**) and 451 natural accession accessions (**b**); absolute growth rate during ontogeny, *GR*(*t*), on one individual (**c**) and the 451 accessions (**d**); relative growth rate during ontogeny, *RGR*(*t*), on one individual (**e**) and the 451 accessions (**f**). *t_inf_* (red dashed line) represents point of growth curve inflection. Black dashed lines in right panels represent 95% CI of fitted growth curve. Individuals on the right panels are coloured by duration (days) of plant life cycle. (**g**-**i**) Variation of *M*(*t*_inf_), *GR*(*t*_inf_) and *RGR*(*t*_inf_) across the 451 accessions phenotyped, with broad-sense heritability (*H*^2^) on the top-left corner of each panel. Dots represent genotypic mean ± standard error (*n* = 2).

Growth dynamics variables were computed from the fitted parameters, such as *GR*(*t*), the derivative of the logistic growth function (Fig. 2c and 2d), as in Equation 2:

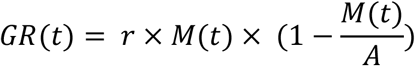

and the relative growth rate (mg d^-1^ g^-1^), *RGR*(*t*), measured as the ratio *GR*(*t*) / *M* (*t*) (Fig. 2e and 2f), as in (Equation 3:

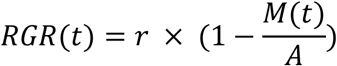

Comparing growth traits measured at *t*_inf_, *i.e.* when *GR* is maximal for all individuals [4], revealed important variation between accessions (Fig. 2g-i), with a high part of phenotypic variance accounted by genetic variability, as measured by broad-sense heritability (*H*^2^ = 0.93, 0.90 and 0.65 for *M*(*t*_inf_), *GR*(*t*_inf_) and *RGR*(*t*_inf_), respectively). To evaluate the robustness of the method, we repeated an experiment on 18 accessions selected for their highly contrasted phenotypes (Fig. S2). Results showed a good correlation between the rosette dry mass at the inflection point estimated in the first experiment and the dry mass measured in the second experiment (*r*^2^ = 0.67; Fig. S3a).

### Estimation of fruit number from inflorescence images

#### Description

The method to estimate fruit number from inflorescence images requires to manually counting fruits on a fraction of individuals in order to train predictive models (Fig. 3). At maturity, inflorescence and rosette of all individuals of the focal population were separated and both photographed with a high-resolution camera (Fig. 3a). Fruits were manually counted on the inflorescence images of 352 out of 856 plants harvested (Fig. 3b). In parallel, we analysed the inflorescence skeletons of all the 856 harvested plants with a dedicated ImageJ macro (Additional File 4). Using the skeleton descriptors computed with the macro and manual measurements in the population subset, we evaluated the accuracy of different models to predict the number of fruit per individual (Fig. 3c), and applied the best model to the whole focal population.

**Figure 3.**
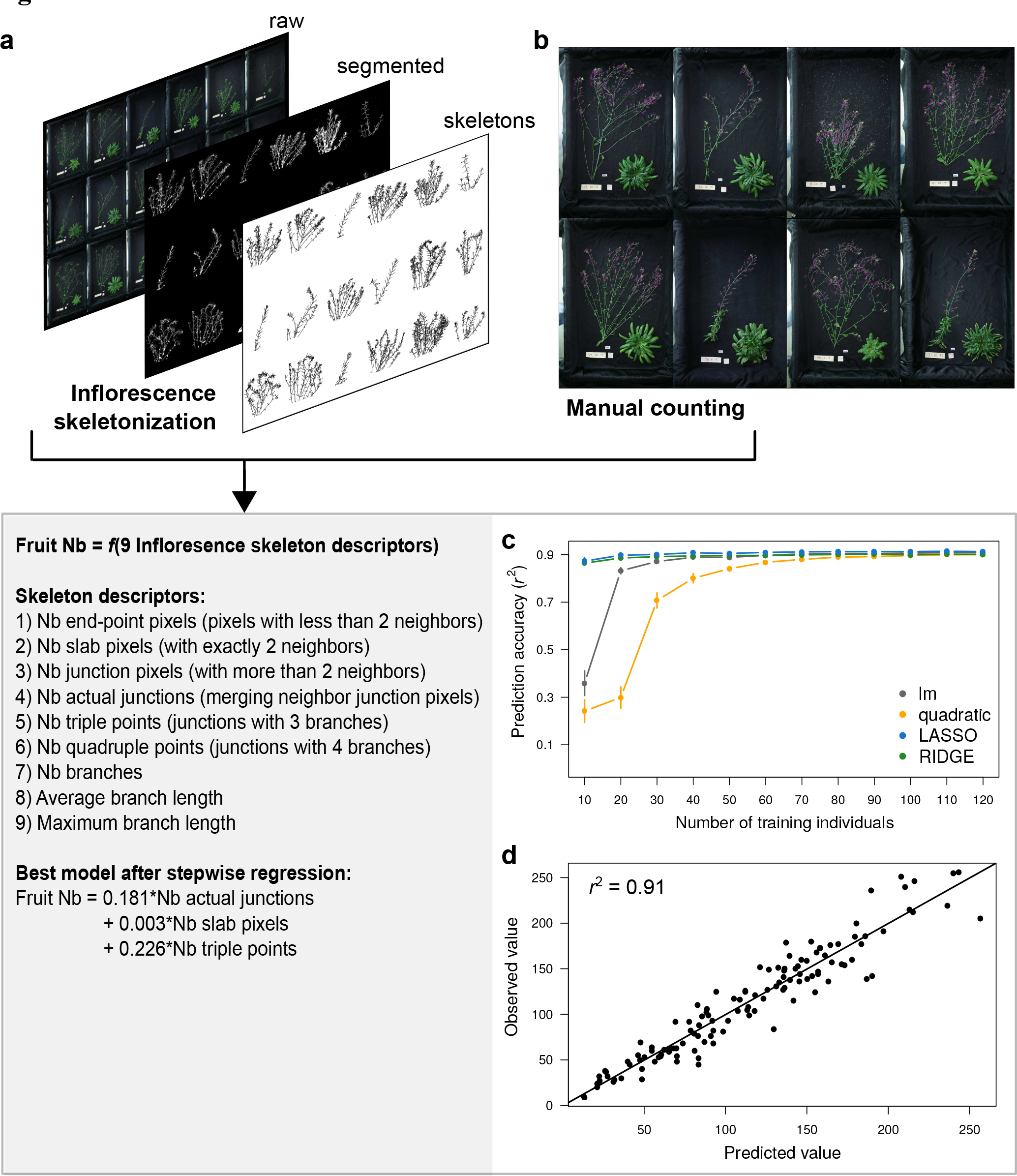
Estimation of fruit number from image analysis and statistical modelling. (**a**) Example of inflorescence images, analyzed with ImageJ to extract individual skeleton descriptors after segmentation and 2D skeletonization. (**b**) Manual counting (purple dots) of fruit number on a subset of inflorescence images. (**c**) Series of cross-validation performed for different predictive models with different training population size (x axis). Dots represent mean prediction accuracy, measured as Pearson’s coefficient of correlation (*r*^2^) between observed and predicted values. Error bars represent 95% CI across 100 random permutations of the training dataset. (**d**) Correlation between observed and predicted values for cross-validation of the best model obtained with stepwise regression, performed 60 individuals to train the model, and tested on 100 individuals not used to train the model.

#### Implementation

After opening ImageJ, users need to go to *Plugins* > *Macros* > *Run*, and select the “RAPAmacro_InflorescenceSkeleton.txt” file in the corresponding folder. Following macro instructions, users have to label the plant and define the zone of interest by drawing a rectangle around inflorescence. For all images present in the input folder, the macro will then automatically perform image segmentation, skeletonization, and computation of 2D skeleton parameters of inflorescence (Fig. 3a). 2D skeletons analysis with ImageJ returns nine vectors of variables for each plant (described in Fig. 3), which were automatically saved as .xls file by the macro (in an output folder defined by user). The sums of these nine vectors per individual were used as nine predictors of fruit number.

Using the same approach as for estimating rosette dry mass, we tested different models and different training population size with cross-validation (Additional File 5). As for rosette dry mass, results showed that the nine skeletons descriptors predict > 90% of fruit number in 100 individuals not used to train the model (Fig 3c). With a training population size > 30 individuals, lm performed equally than LASSO and RIDGE regressions. As for dry mass estimation, quadratic models performed poorly. For small training population size, LASSO and RIDGE regressions reached higher prediction accuracy than linear or quadratic models. Using stepwise regression, we showed that the best model to estimate fruit number with 60 training individuals is *Fruit Nb* = 0.181 × *Nb actual junctions* + 0.003 × *Nb slab pixels* + 0.226 × *Nb triple points*, in a linear model framework (cross-validation *r*^2^ = 0.91, Fig. 3d).

#### Application

The model to estimate fruit number from inflorescence images was applied on all individuals of the focal population (Fig. 4a). We measured a relatively high broad-sense heritability for fruit production across accessions (*H*^2^ = 0.68), compared to *H*^2^ estimates of morphological and physiological traits measured in previous studies [33]. In addition, fruit number estimated from image analysis was well correlated with fruit number manually counted on 18 genotypes phenotyped in a second experiment (*r*^2^ = 0.70; Fig. S3b). To further validate the method, we applied the predictive model on an independent set of inflorescence images taken at the Center for Plant Molecular Biology (ZMBP, University of Tübingen) on the *rev5* knock-out mutant. Compared to wild-type Col-0, *rev5* produces less fruits due to the effect of the mutation on branching pattern and flower development [34]. This was well captured by the predictive model (Fig. 4b), yet trained on the natural accessions.

**Figure 4.**
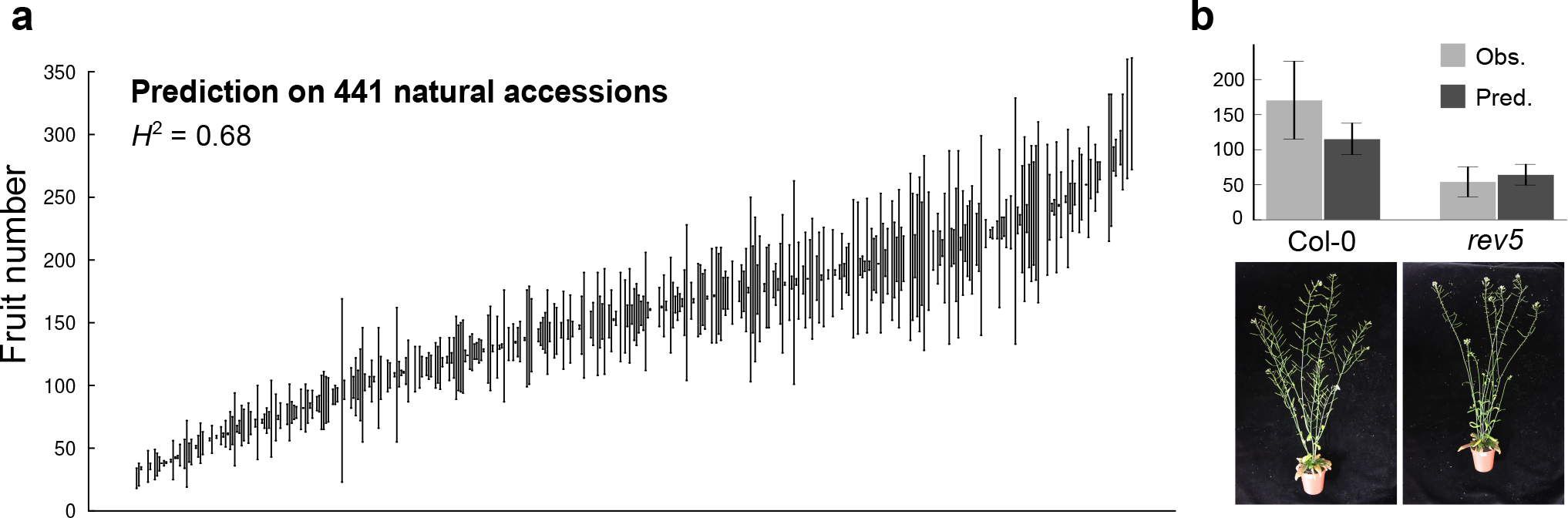
Application of the method to estimate fruit number in natural accessions and *rev5* mutant of *A. thaliana.* (**a**) Variability in fruit number across 441 natural accessions, with broad-sense heritability (*H*^2^) on the top-left corner. Dots represent genotypic mean ± standard error (*n* = 2). (**b**) Prediction of fruit number (mean +/- 95% CI) from model trained on accessions and applied to *rev5* mutant and Col-0 wild-type (*n* = 5). Results are compared to observed fruit number manually counted at harvesting.

## Discussion

RGR and fruit production are two key physiological parameters related to individual performance and fitness [1–3], but they remain difficult to measure in large scale experiments. Our goal was to develop a set of tools for biologists to analyze these traits with low-cost equipment. From simple top-view imaging of rosette and inflorescence, we built robust predictive models of plant dry mass and fruit number. Based on a semi-invasive approach and two computing steps - one to analyze images with ImageJ and one to model data with R -, the method allows a single experimenter to simultaneously measure biomass accumulation, RGR and fruit production over thousands of plants.

Top view pot or tray imaging can easily be done in any laboratory or facilities. In this study, we used pictures of trays manually taken during ontogeny with a regular camera. The same approach has been proposed in low-cost systems for high-throughput phenotyping in *A. thaliana*, using projected rosette area to measure growth during several hours or days [14–16]. Comparatively, our method allows measuring the absolute and relative rate of biomass accumulation during the whole life cycle of a plant. The time lapse and frequency of tray imaging is important for proper fitting of the growth curve. We used daily imaging during the 25 first days of growth after vernalization, although growth curves can be fitted with only one picture every 2-3 days. The ImageJ macro we developed here automatically processes tray images when plants are young and do not overlap. When they become too large (20-25 DAG in our study), the macro offers the possibility to spatially separate plants (in manual mode). We estimated that, on a desktop computer, the macro takes approximately 20-25 s per tray (30 individuals) when running on automatic mode, and between one and two minutes in manual mode (depending on the number and amplitude of corrections to make).

The semi-invasive approach drastically reduces the number of replicates necessary to measure the growth dynamics, or the time needed for manual measurement of fruit number. Furthermore, it allows experimenter to compute biomass accumulation non-destructively until maturity, and thus, to compare growth and reproductive success on the same individuals. We showed that the method is robust and reproducible across experiments when performed in the same environment. Moreover, the model for fruit prediction correctly predicted the decrease in *rev5* in a complete independent experiment. However, we recommend making a new predictive model of plant biomass with cross-validation for each experiment (code example available in Additional File 2), specifically if growth conditions change, as the relationship between rosette morphology and rosette biomass is expected to differ depending on genotypes and environments.

In this study, we propose flexible methods and customizable tools for researchers to characterize plant phenotype in their own facilities. This should lower the barriers of high-throughput phenotyping, and help dissect the relationships between growth dynamics and reproductive success in various laboratory conditions. Methods were developed for *A. thaliana*, which is the favourite model in plant genetics and molecular biology, and it is also becoming a model in evolutionary biology and ecology [35–37]. The same approach is nonetheless applicable to other species, although 3D architecture requires more sophisticated image analysis. A recent study in maize offers a nice example of 3D reconstruction and biomass prediction with a dedicated phenotyping platform [8].

## Methods

### Plant material

472 natural accessions of *A. thaliana* were selected from the initial germplasm of the 1001 Genomes project [29] (http://1001genomes.org/; Table S1). Seeds used in this study were obtained from parental plants propagated under similar conditions in greenhouse. All seeds were stored overnight at -80 °C and surface-sterilized with 100% ethanol before sowing. A transgenic line of *A. thaliana* affecting in branching pattern and fruit production was used: *rev5*, which has a strong ethyl-methylsulfonate (A260V) knock-out mutation of *REVOLUTA* in the Columbia Col-0 background [34].

### Growth conditions

Plants were cultivated in hydroponics, on inorganic solid media (rockwool cubes) watered with nutrient solution [38]. Four replicates of 472 accessions were grown, with pots randomly distributed in 64 trays of 30 pots each. Seeds were sowed on 3.6 cm x 3.6 cm x 4 cm depth rockwool cubes (Grodan cubes, Rockwool International A/S, Denmark) fitted in 4.6 cm (diameter) x 5 cm (depth) circular pots (Pöppelmann GmbH and Co., Germany). The pots were covered with a black circle pierced in the centre (5-10 mm hole manually made with a puncher). Before sowing, the dry rockwool cubes were watered with 75% strength nutrient solution. The chemical composition of the nutrient solution was obtained from Conn *et al.* [38].

After sowing, trays were incubated for two days in the dark at 4 °C for seed stratification, and then transferred for six days to 23 °C for germination. After germination, all plants, having two cotyledons, were vernalized at 4 °C during 41 days to maximize the flowering of all the different accessions. Plants were thinned to one plant per pot, and trays were moved to the RAPA room, set to 16 °C with a temperature variability of close to ± 0.1 °C, air humidity at 65 %, and 12 h day length, with a PPFD of 125 to 175 µmol m^-2^ s^-1^ provided by a 1:1 mixture of Cool White and Gro-Lux Wide Spectrum fluorescent lights (Luxline plus F36W/840, Sylvania, Germany). All trays were randomly positioned in the room, and watered every day with 100% strength nutrient solution.

Replicates 1 and 2 (the focal population, *n* = 960) were harvested at maturity, when fruit senescence started. Due to germination failure and mortality, only 451 accessions were phenotyped for growth, and 441 for fruit number. Replicates 3 and 4 (the training population, *n* = 960) were harvested at 16 DAG for dry mass measurement.

A second experiment was performed on a set of 18 contrasted accessions (Fig. S2), grown in the same conditions. Three replicates per genotype were harvested at the estimated inflection point for rosette dry mass measurement (inflection point estimated from the first experiment), and five replicates were harvested at the end of the life cycle for manual fruit counting.

*rev5* and Col-0 were cultivated in the Center for Plant Molecular Biology (ZMBP, University of Tübingen, Germany). Plants were grown on standard soil (9:1 soil and sand) under controlled conditions: in long days (16 h day; 8 h night), low light (70-80 μE m^-2^ s^-1^) and an ambient temperature of 21 °C (see [39] for details).

### Plant imaging and harvesting

All trays were manually imaged every day during the first 25 days after vernalization with a high-resolution camera (Canon EOS-1, Canon Inc., Japan). Individual labelling (*i.e.* genotype, replicate and date of measurement) was performed with ImageJ [31] during the image analysis process with the “RAPAmacro_RosetteShape.txt” macro. Image segmentation was used to separate background from plant. The same segmentation was performed on rosette and inflorescence, after inverting images and adjusting color threshold, by setting hue between 178 and 255, brightness between 178 and 255, and saturation between 35 and 255. However, it is important to note that color threshold for segmentation depends on light conditions during imaging, and thus, need to be adjusted by users on a set of test images. Undesirable dots that remained after segmentation were removed with the ‘Remove outliers’ option in ImageJ to clean segmented images. After segmentation, inflorescence skeletonization and 2D skeleton analysis were automatically performed with the corresponding functions in ImageJ (see code in Additional File 4). Skeletons were not pruned for loops. Extracted rosette shape and inflorescence skeleton parameters were automatically saved in .xls format.

Plants in the training population were harvested at 16 days after vernalization, rosette were dried at 65 °C for three days, and separately weighed with a microbalance (XA52/2X, A. Rauch GmbH, Graz, Austria). All individual rosette parameters extracted after segmentation were saved as .xls files, each row corresponding to a specific date, tray label and pot coordinates.

At fruit maturity, inflorescence and rosette of the focal population were harvested and separately photographed. Rosettes were dried at 65 °C for at least three days, and weighed with a microbalance (XA52/2X). In the experiment at ZMBP, whole plants of *rev5* and Col-0 were photographed at fruit maturity (first yellowish fruits) by taking side pictures of each pot separately (n = 5).

### Statistical analyses

Different predictive models were evaluated for both the estimation of rosette dry mass and fruit number. We notably compared linear models, quadratic models - where each predictor was fitted as a two-order polynomial function -, RIDGE and LASSO regressions (Additional Files). Prediction accuracy was tested by cross-validation on 100 individuals not used to train the model, using the Pearson’s coefficient of correlation (*r*^2^) between observed and predicted trait values. For each model, we tested prediction accuracy according to training population size across 100 random permutations of the training dataset (Additional Files). Training population size varied between 10 and 250 for dry mass estimation, and between 10 and 120 for fruit number estimation. Stepwise regression, using *step* function in R, was used to identify the best model, with minimum predictors, of rosette dry mass and fruit number.

Non-linear fitting of individual growth curves (Equation 1) were performed with the *nls* function in R (Additional File 3). Since some plants germinated during or, for a few, after vernalization, we considered the first day of growth (*t*_0_) for each individual of the focal population as the day at which it had a minimum size. For convenience, we used the size of the biggest measured plant across all individuals at the first day of growth following vernalization, which corresponded to a plant with first true leaves just emerged. Growth was expressed as a function of days after germination (DAG, starting at *t*_0_). This procedure allowed for normalization of growth trajectories from the same starting point between individuals that differ in germination speed [40]. Growth dynamics variables were computed from the fitted parameters (mean and 95% confidence interval (CI)), such as absolute growth rate, *GR*(*t*), the derivative of the logistic growth function (Equation 2), and *RGR*(*t*) (Equation 3).

Broad-sense heritability (*H*^2^) was calculated with a Bayesian approach implemented in a *MCMCglmm* model performed in R, considering the accession as a random factor, as:

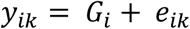

where y is trait of interest in individual *k* of genotype *i*, *G_i_* is the accession *i*, and *e_ik_* is the residual error. *H*^2^ was calculated at the proportion of genotypic variance (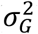) over total variance (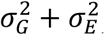):

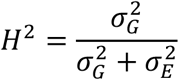

## Declarations

### List of abbreviations

*t*_0_: first day of growth after vernalization
*t*_inf_: inflection point (days) of the logistic growth curve
*A*: upper asymptote of the logistic growth curve (mg)
*B*: inverse of the exponential constant of the logistic growth curve
DAG: days after *t*_0_
M: rosette dry mass (mg)
GR: absolute growth rate (mg d^-1^)
RGR: relative growth rate (mg d^-1^ g^-1^)

### Ethics approval and consent to participate

Not applicable.

### Consent for publication

Not applicable.

### Availability of data and material

The datasets supporting the conclusions of this article are included within the article and its additional files. Codes are available on Github (https://github.com/fvasseur), and phenotypic data are available on the Dryad repository (http://datadryad.org/) [41]. Correspondence and requests for materials should be addressed to.

### Author Contributions

FV, GW, JB, RS and DW designed the study. FV, RS and GW performed the experiments and extracted the data. FV, GW and JB performed image and statistical analyses, and wrote codes. FV interpreted the results and wrote the paper.

## Acknowledgements

We thank Jane Devos for her help in performing the experiments, and Denis Vile, Dominik Grimm and Moises Exposito-Alonso for helpful comments about image analysis and statistical modelling. We also thank Ulrike Zentgraf and Stephan Wenkel for providing material and data to assess fruit number in *rev5* mutant.

## Additional Files

**Additional File 1. “RAPAmacro_RosetteShape.txt”.** ImageJ macro used to extract rosette shape descriptors from top-view tray or pot images.

**Additional File 2. “RAPAcode_EstimationDryMass.R”.** R code used to predict rosette dry mass from rosette shape descriptors, with cross-validation approach to train and test different models and training population size.

**Additional File 3. “RAPAcode_ModellingGrowthDynamics.R”.** R code used to model sigmoid growth curves and growth dynamics (*M*(*t*), *GR*(*t*), and *RGR*(*t*)) from predicted rosette dry mass during ontogeny and measured rosette dry mass at maturity.

**Additional File 4. “RAPAmacro_InflorescenceSkeleton.txt”.** ImageJ macro used to extract inflorescence skeleton descriptors from top-view images of plant inflorescence.

**Additional File 5. “RAPAcode_EstimationFruitNumber.R”.** R code used to predict fruit number from inflorescence skeleton descriptors, with cross-validation approach to train and test different models and training population size.

**Additional Figure S1. The RAPA facility.** Entrance view of the growth chamber with a zoom on the camera installed between light tubes (top-left panel). On the right is the setup to water the plants and take manual tray pictures.

**Additional Figure S2. Representation of the 18 accessions phenotyped in the second experiment.** Nine phenotypic groups represented by the purple circles (three groups of RGR and three groups of growth duration) were selected, each containing two accessions.

**Additional Figure S3. Inter-experiment reproducibility of rosette dry mass and fruit number estimation.** Measured across 18 contrasted accessions. (**a**) Pearson’s coefficient of correlation (*r*^2^) between rosette dry mass *M* estimated at the inflection point *t*_inf_ in the first experiment and rosette dry mass *M* measured at *t*_inf_ in the second experiment. (**b**) *r*^2^ between the number of fruits estimated in the first experiment and the number of fruits measured in the second experiment.

**Additional Table S1.** List of the 451 accessions phenotyped (*n* = 2), with measured and estimated traits, and fitted model parameters of Equation 1.

